# Nanobodies for the treatment of SARS-CoV-2 in animals: a meta-analysis

**DOI:** 10.1101/2022.09.26.509459

**Authors:** Jiao Jiao, Shulan Lin, Zhenqi Liang, Peng Wu

**Affiliations:** College of life Sciences, Shihezi University, Shihezi, Xinjiang, China

**Author notes:** Corresponding author: Peng Wu, College of life Sciences, Shihezi University, China.

**Keywords:** COVID-19, SARS-CoV-2, animal, VHHs, treatment, meta-analysis

## Abstract

This meta-analysis aimed to find the effect of variable domain of heavy-chain antibodies (VHHs) for the treatment of severe acute respiratory syndrome coronavirus-2 (SARS-CoV-2) in animals. The databases of the PubMed, China National Knowledge Infrastructure (CNKI), Wan fang data, Cochrane Library, and Embase were searched for articles published before August 2022 on the protective effects of VHHs in animals. The articles retrieved were screened using inclusion and exclusion criteria. The data were analyzed using Review Manager 5.4. Six articles were selected from 667 articles based on the inclusion and exclusion criteria in VHHs. A forest plot showed that VHHs could offer protection against SARS-CoV-2 infection in animals [Mantel-Haenszel (MH) = 172.94, 95% confidence interval (CI) = (43.96, 678.42), P < 0.00001]. There was almost no heterogeneity in this study (I^2^ = 0). A funnel plot showed that the bias of the data analysis was small. This is a special meta-analysis proved that VHHs could treat and prevent SARS-CoV-2 in animals.

## Introduction

Coronavirus disease 2019 (COVID-19) is a highly contagious disease caused by severe acute respiratory syndrome coronavirus-2 (SARS-CoV-2). The COVID-19 pandemic has become a serious burden on global public health ^[1]^. Although the world has made unprecedented efforts to rapidly develop a vaccine for SARS-CoV-2, the situation is still grim ^[2]^. The Omicron variant has an unprecedented devastating effect on immune protection established by vaccination and infection ^[3]^.

Therapeutic neutralizing antibodies are considered to be an effective method for treating COVID-19 in the short and medium terms ^[4]^. The pandemic of COVID-19 highlights the serious consequences of animal virus leakage on public health, economy and society ^[5]^. A worrying problem is that SARS-COV-2 may spread to local species, which may make the virus popular by setting up a second host ^[6]^. The widespread of SARS-COV-2 in human beings increases the theoretical risk of reversal of zoonotic events in animals, that is, SARS-CoV-2 is introduced into animals^[7]^.

VHH is the smallest functional antigen-binding fragment found in camel blood ^[8]^. VHH cocktails have further provided effective therapeutic activity against the Omicron variant infections in hamsters ^[3]^. VHHs have great potential for the treatment of COVID-19. In this meta-analysis, the research status of VHH therapy for COVID-19 was summarized and evaluated in animals.

## 1 Methods

### 1.1 Reporting

This meta-analysis was conducted according to the preferred reporting items of the systemic review and meta-analysis guidelines (S1 supporting information form).

### 1.2 Literature search strategy

The databases of the CNKI, PubMed, Wan fang data, Cochrane Library, and Embase were searched for articles published before August 2022 by PW and JJ. “SARS-CoV-2” or “COVID-19” and “VHH” were used as the search keywords.

### 1.3 Inclusion and exclusion criteria

Inclusion criteria: ➀ Published Chinese or English articles on SARS-CoV-2 VHHs. ➁ The same article contains animal experiments. ➃ The article must comprise a challenge of SARS-CoV-2. ➄ The antibodies used in the article comprise a VHH, nanobody, and a single domain heavy chain antibody.

Exclusion criteria are as follows: ➀ review articles; κ articles without animal or clinical experiments; ➁ duplicate articles or studies; and ➃ articles without SARS-CoV-2 experiments.

### 1.4 Data extraction

PW and JJ initially screened the articles by reading their titles. Then, using the exclusion criteria, we read the abstract and the full text and performed independent data extraction. The number of protected animals and the total number of animals were extracted as the main data. Different opinions were resolved through discussions. We performed a subgroup analysis of the protective effects on hamsters and mice.

### 1.5 Statistical analysis

Data analysis was performed using Review Manager 5.4 (Cochrane collaboration). In the articles, I^2^ was used to evaluate the heterogeneity: when I^2^ < 50%, the fixed effect model was selected for meta-analysis, and the random effect model was selected when I^2^ > 50%. Odds ratio (OR) with 95% confidence intervals was applied for the dichotomous variables. The Mantel-Haenszel method was used to verify the data. A funnel graph was used to judge the publication bias of the extracted data.

## 2 Results

### 2.1 Literature screening results

Based on the retrieval scheme, 667 articles were retrieved, of which 95 were duplicate articles. After reading the titles and abstracts based on the exclusion criteria, 48 articles were selected. Finally, six articles fully met our inclusion criteria (Fig 1).

**Figure 1.**
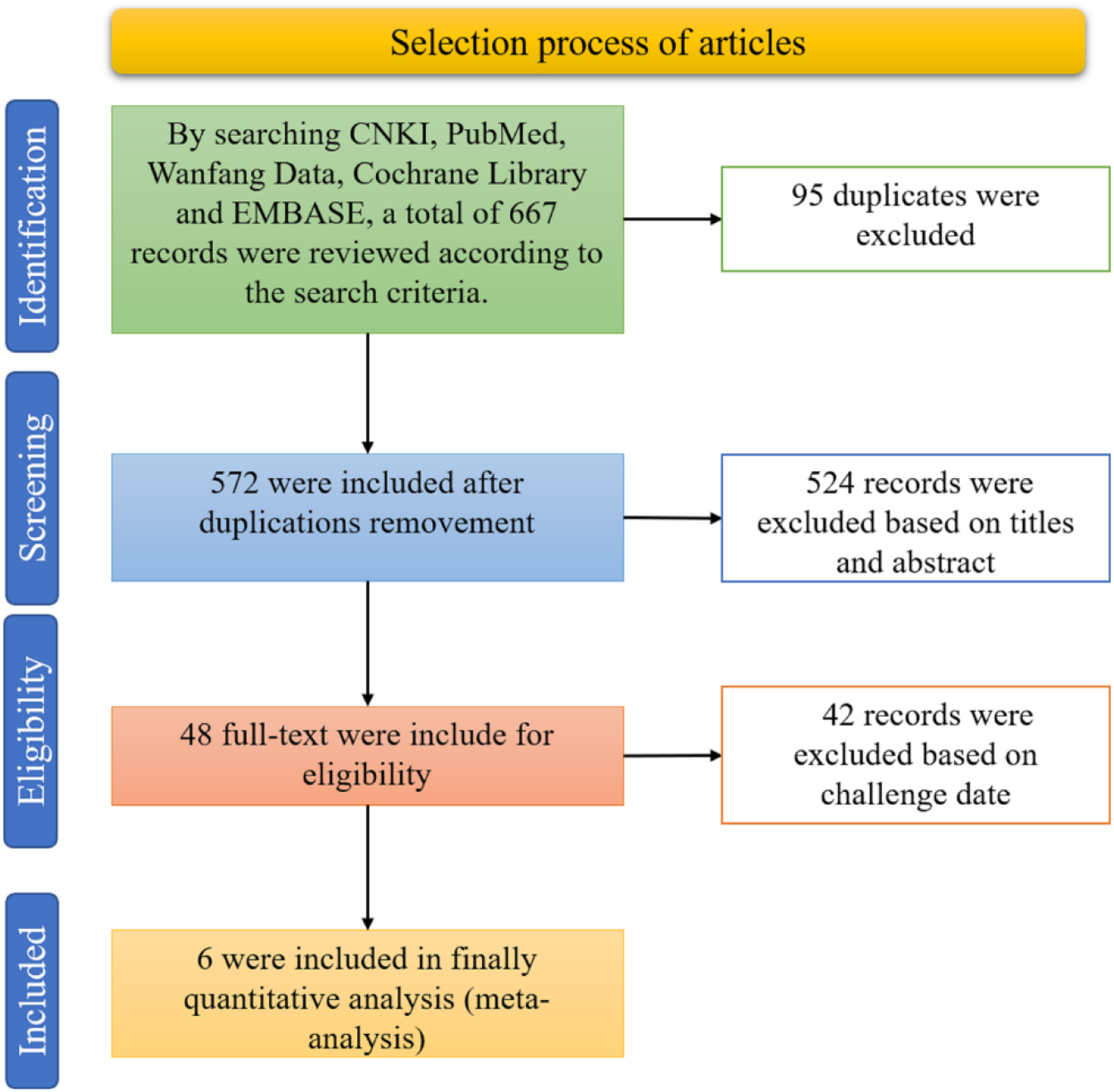
PRISMA flow diagram of search results

### 2.2 Experimental data extraction

The data extracted are summarized in Table 1, comprising a total of 180 animals. The articles were published between 2020 and 2022. The strains were SARS-CoV-2, Omicron BA.1 variant, SARS-CoV-2 Beta variant, and hCoV-9/Australia/VIC2089/2020. The VHHs used were administered through intraperitoneal injection, aerosol spray, and intranasal injection. The amount of VHHs administered ranged from 0.2–20 mg/kg. The dose for the challenge test ranged from 1×10^4^-1×10^6^ cfu. The methods used for the poison attack experiments were intranasal and glas-col nebulization inhalation.

**Table 1.** Characteristics and summary findings of the selected studies

|  | Author | Year | Control Group |  | Experimental Group |  | Strain | Approche | Nanobody Dose | Challenge Dose (pfu) | Challenge Approche |
| --- | --- | --- | --- | --- | --- | --- | --- | --- | --- | --- | --- |
|  |  |  | Protected Number | Total Number | Protected Number | Total Number |  |  |  |  |  |
| 1 | Gang Ye | 2020 | 0 | 6 | 18 | 18 | SARS-CoV-2 | Intraperitoneally | 10-20 mg/kg | 1000000 | Intranasally |
| 2 | Huan Ma-A | 2022 | 0 | 5 | 5 | 5 | SARS-CoV-2 | Intraperitoneally | 10 mg/kg | 20000 | Intranasally |
| 3 | Huan Ma-B | 2022 | 0 | 6 | 10 | 10 | Omicron BA.1 variant | Intraperitoneally | 10 mg/kg | 10000 | Intranasally |
| 4 | Phillip Pymm | 2021 | 0 | 10 | 40 | 45 | hCoV-19/Australia/VIC2089/2020 | Intraperitoneally | 0.2-5 mg/kg | 15000000 | Glas-col nebulization inhalation |
| 5 | Sham Nambulli | 2021 | 0 | 6 | 5 | 6 | SARS-CoV-2 | Aerosolization | None | 30000 | Intranasally |
| 6 | Siling Wang-A | 2022 | 0 | 5 | 15 | 15 | SARS-CoV-2 Beta variant | Intraperitoneally | 20 mg/kg | 10000 | Intranasally |
| 7 | Siling Wang-B | 2022 | 0 | 6 | 6 | 6 | SARS-CoV-2 Omicron variant | Intraperitoneally | 20 mg/kg | 10000 | Intranasally |
| 8 | Xilin Wu | 2021 | 0 | 4 | 27 | 27 | SARS-CoV-2 | Intranasally | 10 mg/kg | 100000 | Intranasally |

### 2.3 Data synthesis

The results of the forest plot show that VHHs could offer protection against SARS-CoV-2 infection in animals [MH =172.94, 95% CI = (43.96, 678.42), P < 0.00001] (Fig 2). The sensitivity is acceptable (df = 7 (P = 0.99)). The sensitivity is acceptable as I^2^ = 0%.

**Figure 2.**
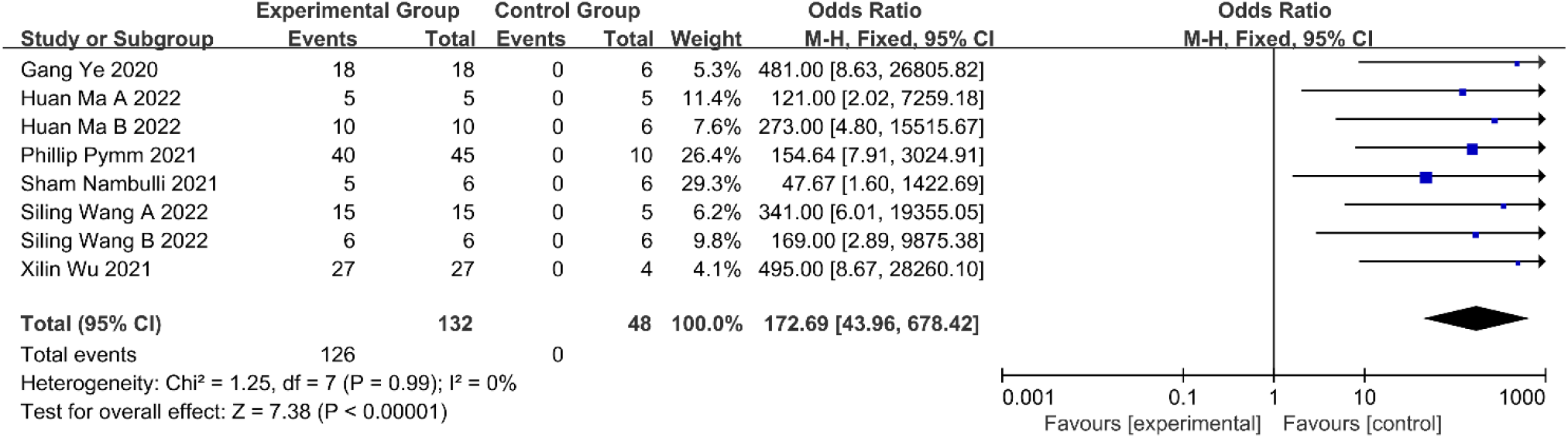
Forest plot of the ability of VHHs to protect animals.

The results of the forest plot show that VHHs could offer protection against SARS-CoV-2 infection in hamsters [MH = 168.05, 95% CI = (28.44, 992.86), P < 0.00001] (Fig 3). The sensitivity is acceptable (df = 4 (P = 0.92)). The sensitivity is acceptable as I^2^ = 0%.

**Figure 3.**
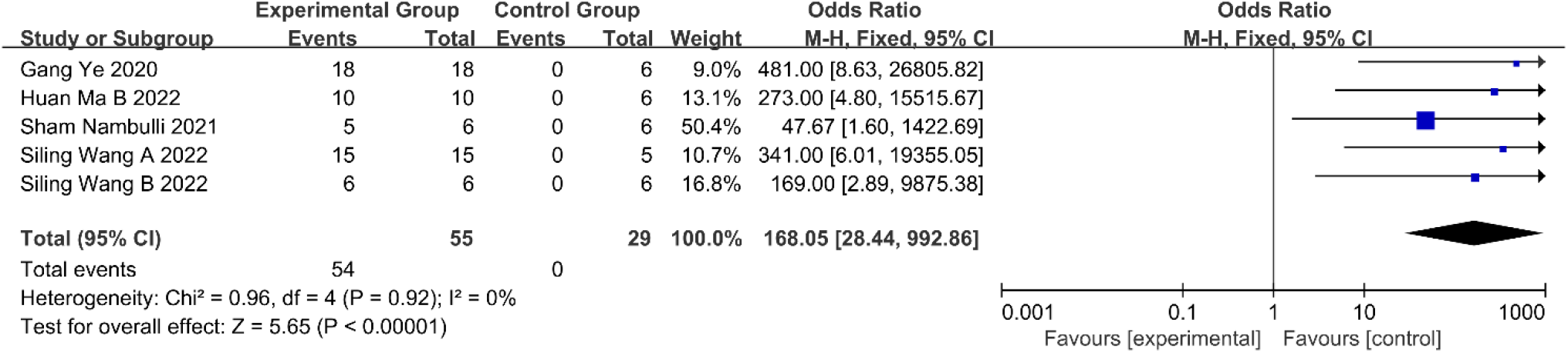
Forest plot of the ability of VHHs to protect hamsters.

The results of the forest plot show that VHHs could offer protection against SARS-CoV-2 infection in mice [MH = 179.13, 95% CI = (21.06, 1523.36), P < 0.00001] (Fig 4). Three articles on the ability of VHHs to protect mice were selected.

**Figure 4.**
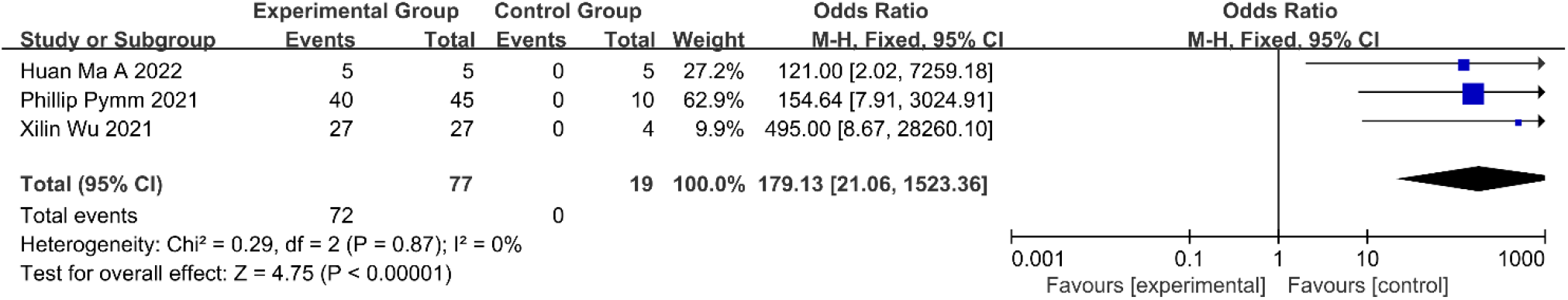
Forest plot of the ability of VHHs to protect mice.

### 2.4 Publication bias

Note: (A) Funnel plots of the ability of VHHs to protect animals, (B) Funnel plots of the ability of VHHs to protect hamsters, and (C) Funnel plots of the ability of VHHs to protect mice. Table 1. Characteristics and summary findings of the selected studies

The publication bias is small because the funnel graph is symmetrical (Fig 5). This shows that the experimental data of the articles we selected were rigorous.

**Figure 5.**
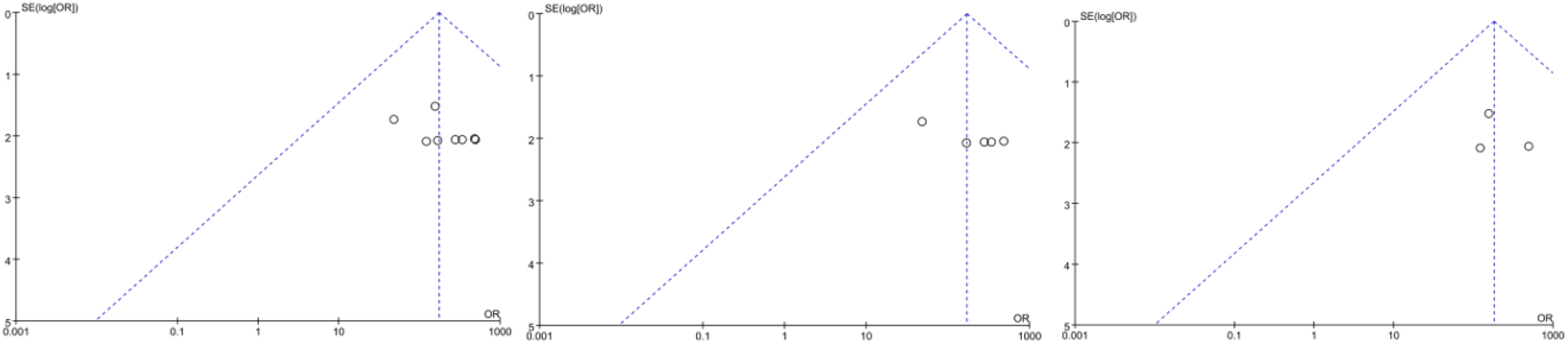
Funnel plots.

## 3 Discussion

Because of their small size, simple and robust structure, high thermal stability, and good solubility, VHHs are delivered directly to the lungs via an inhaler ^[9]^. VHHs are highly stable and well suited for development as biological inhalation therapies for respiratory diseases ^[10]^. Ablynx has developed an inhaled antirespiartory syncytial virus VHH, alx-0171, with potent antiviral effects and reduced viral infection symptoms in animal models ^[4]^. A key requirement for pulmonary drug delivery is the stability of the biological product to allow it to withstand the degradative environment ^[11]^. An efficient bispecific VHH protected hACE2 mice from SARS-CoV-2 infection through intranasal administration ^[12]^.

VHHs have many advantages in virus therapy. The complementarity detecting region CDR-3 of VHHs are relatively long, which can increase the affinity for antigen binding, and VHHs can bind to places where traditional antibodies cannot ^[13]^. Using bioengineering to connect VHHs with albumins and other molecules with long half-life can slow down the clearance time of VHHs, which can achieve a better therapeutic effect ^[14]^.

Monoclonal antibodies cannot enter tissues, which limits the use of monoclonal antibodies ^[15]^. A VHH is only one-tenth the size of a monoclonal antibody, and it is the smallest natural fragment ^[16]^. VHHs have the advantages of high solubility, high domain stability, and the ability to cross the blood-brain barrier ^[17]^. VHHs have advantages over monoclonal antibodies in many aspects. VHHs can be expressed in large quantities by prokaryotic expression systems. Using VHHs instead of monoclonal antibodies can reduce the cost of treatment ^[18]^.

There are many limitations to this research. Different doses of VHHs may lead to deviations. Studies in other languages that were not selected may have also led to the appearance of bias. It is possible that some studies with poor results have not been published in journals. Currently, many studies have shown that VHHs can neutralize SARS-CoV-2 in vitro, but only six articles have been verified in vivo. The published articles were limited to hamsters and mice.

## 4 Conclusions

This meta-analysis proved that VHHs could treat and prevent SARS-CoV-2 in animals. Before they can be used, further preclinical analysis is needed, including extensive toxicological pathology research.

## Notes

### Competing Interest Statement

The authors have declared no competing interest.

